# Modulation of Grasp Parameters using Arbitrary Cues about Object Property in Older Adults

**DOI:** 10.1101/2020.10.19.344457

**Authors:** Nishant Rao, Neha Mehta, Pujan Patel, Pranav J. Parikh

**Author notes:** **Corresponding Author:** Pranav J. Parikh, Ph.D. Department of Health and Human Performance 3875 Holman Street, suite 104R GAR University of Houston Houston, TX 77204, USA.

## Abstract

Dexterous manipulation may be guided by explicit information about object property. Such a manipulation requires fine modulation of digit position and forces using explicit cues. Young adults can form arbitrary cue-object property associations for accurate modulation of digit position and forces. Aging, in contrast, might alter this conditional learning. Older adults are impaired in accurately modulating their digit forces using explicit cues about object property. However, it is not known whether older adults can use explicit cues about object property to modulate digit position. In this study, we instructed ten healthy older and ten young adults to learn a manipulation task using arbitrary color cues about object center of mass location. Subjects were required to exert clockwise, counterclockwise, or no torque on the object according to the color cue and lift the object while minimizing its tilt across sixty trials. Older adults produced larger torque error during the conditional learning trials than young adults. This resulted in a significantly slower rate of learning in older adults. Older, but not young adults, failed to modulate their digit position and forces using the color cues. Similar aging-related differences were not observed while learning the task using implicit knowledge about object property. Our findings suggest that aging impairs the ability to use explicit cues about object property to modulate both digit position and forces for dexterous manipulation. We discuss our findings in relation to age-related changes in the processes and the neural network for conditional learning.

## INTRODUCTION

We can perform dexterous manipulation based on explicit information about object properties (1, 5, 7, 16, 24, 27, 28, 32). For instance, subjects can accurately scale digit forces in an anticipatory manner based on arbitrary cues about object weight and texture within a few trials and this associative memory for digit force scaling would last for at least 24 hours (28). Interestingly, when subjects were asked to use visual instructional cues about object CM location to lift the object by preventing it from rolling, they failed to scale their digit forces leading to task failure (36). These findings suggest that such explicit cues of object center of mass location (CM) are ineffective in guiding manipulation that primarily depends on accurate scaling of digit forces. Importantly, dexterous manipulation may also depend on the modulation of digit position based on object properties (9, 13, 25, 31). In the aforementioned association studies, subjects were directed to grasp at the same points on the object, which restricted a change in digit position from trial to trial. That is, an important component of dexterous manipulation – choice of digit position, was neglected. Remarkably, when subjects were given a choice regarding where to grasp the object, they were able to learn to use arbitrary color cues about object CM location to modulate their digit position and digit forces to lift the object while minimizing its tilt (32).

Aging might influence the ability to modulate grasp parameters using arbitrary visual cues about object property for dexterous manipulation (7). Cole and Rotella (7) instructed older adults to grip the object approximately at the center of contact surfaces and lift it using arbitrary cues about either object weight or texture. The authors found that older adults, unlike young adults, showed impaired learning to scale their digit forces using the arbitrary visual cues informing them about object weight and texture preceding each trial (7). Now, the question is whether allowing choice of digit placement would afford older adults an ability to use arbitrary visual cues about object property. Our previous study showed that despite facing an initial difficulty in alignment of digit position while performing dexterous manipulation based on implicit knowledge of a symmetrically shaped object, older adults corrected the digit misplacement within a few lifts due likely to somatosensory and visual information available during task performance (30). It remains unknown whether older adults will be able to modulate their digit positions based on arbitrary visual cues about object property.

The aim of this study was to determine whether older adults can modulate their digit positions based on arbitrary color cues about object CM location for successful object manipulation. We used our recently developed novel conditional visuomotor task that requires subjects to associate an arbitrary color cue to the mass distribution of an object (32). On each trial, subjects had to select one of three possible responses (i.e., exertion of clockwise, counterclockwise, or no torque) according to the color cue and to lift the object while minimizing object tilt across 60 trials. The direction and magnitude of object tilt during the lift provided feedback to subjects about success or failure of their association. We hypothesized that older adults will be able to learn to modulate their digit position based on arbitrary color cues about object CM location. However, as noted earlier (7), older adults will not be able to scale their digit forces in anticipatory manner based on the available arbitrary visual cues. Anticipatory modulation of both digit forces and digit position based on the arbitrary color cues about object CM is necessary to achieve the instructed task goal (32). Therefore, a lack of modulation of digit position and digit forces will result in a failure to accurately scale the compensatory torque based on color-CM association until the time of object liftoff (i.e. anticipatory) in older adults when compared with young adults.

## MATERIALS AND METHODS

### Participants

Twenty participants were recruited across two groups: older adults (69 ±5 yr [mean ±SD], n=10, 4 females) and young adults (25 ±3 yr [mean ±SD], n=10, 5 females). All participants provided written informed consent to participate in the study. Participants had normal or corrected-to-normal vision and no self-reported history of neurological disease and musculoskeletal disorders or upper limb injury. All participants followed the same experimental protocol. Tactile sensibility thresholds were obtained from the distal volar pads of the index finger using Semmes-Weinstein pressure filaments (Smith and Nephew Roland, Menominee Falls, WI). We used a descending method of limits to establish a threshold (30, 34). The index finger was tested approximately midway between the center of the pad and the radial margin of the finger. A threshold was recorded for the smallest filament diameter (buckling that could be perceived on at least 70% of its applications). Both groups demonstrated normal for age tactile sensibility threshold measured using Semmes-Weinstein pressure filaments. The mean tactile sensibility threshold was 208 mg for older adults and 52 mg for young adults. The study was approved by the Institutional Review Board at the University of Houston.

### Apparatus

#### Grip device

A custom-designed inverted T-shaped device was outfitted with two six-dimensional force and torque transducers (**Fig. 1A;** Nano-25; ATI Industrial Automation, Garner, NC) (32, 35). The surfaces of the grip device were covered with sandpaper (grit #320). This device measured grip and load forces (normal and tangential to the graspable surface) and the center of pressure of both thumb and index finger. The base of our grip device consisted of three compartments (left, center, and right). A 400-g mass was inserted in the left (LCM) or right (RCM) bottom compartment to shift the mass distribution of the grip device to the left or right of its vertical midline, respectively. This added mass to either the left or right base compartment generated a negative or positive external torque of 255 N·mm on the device, respectively, following the lift onset. When the added mass was placed in the center base compartment (CCM), the mass distribution remained symmetrical. To prevent visual identification of the location of the added mass, the view of the base compartments was blocked by a lid. As an additional preventive measure, the base compartments were placed away from the participant. The total mass of the grip device with added mass was 790 g. The subject’s hand rested on a table that was placed directly in front of their seat to ensure that the start position and arm posture were consistent throughout the experiment. An electromagnetic sensor (Polhemus FASTRAK; 0.05° resolution) was attached to the top of the device to measure the object roll. The roll of the object was defined as the angle between the gravitational vector and the vertical axis of the grip device, contained in the frontal plane of the grip device.

**Figure 1.**
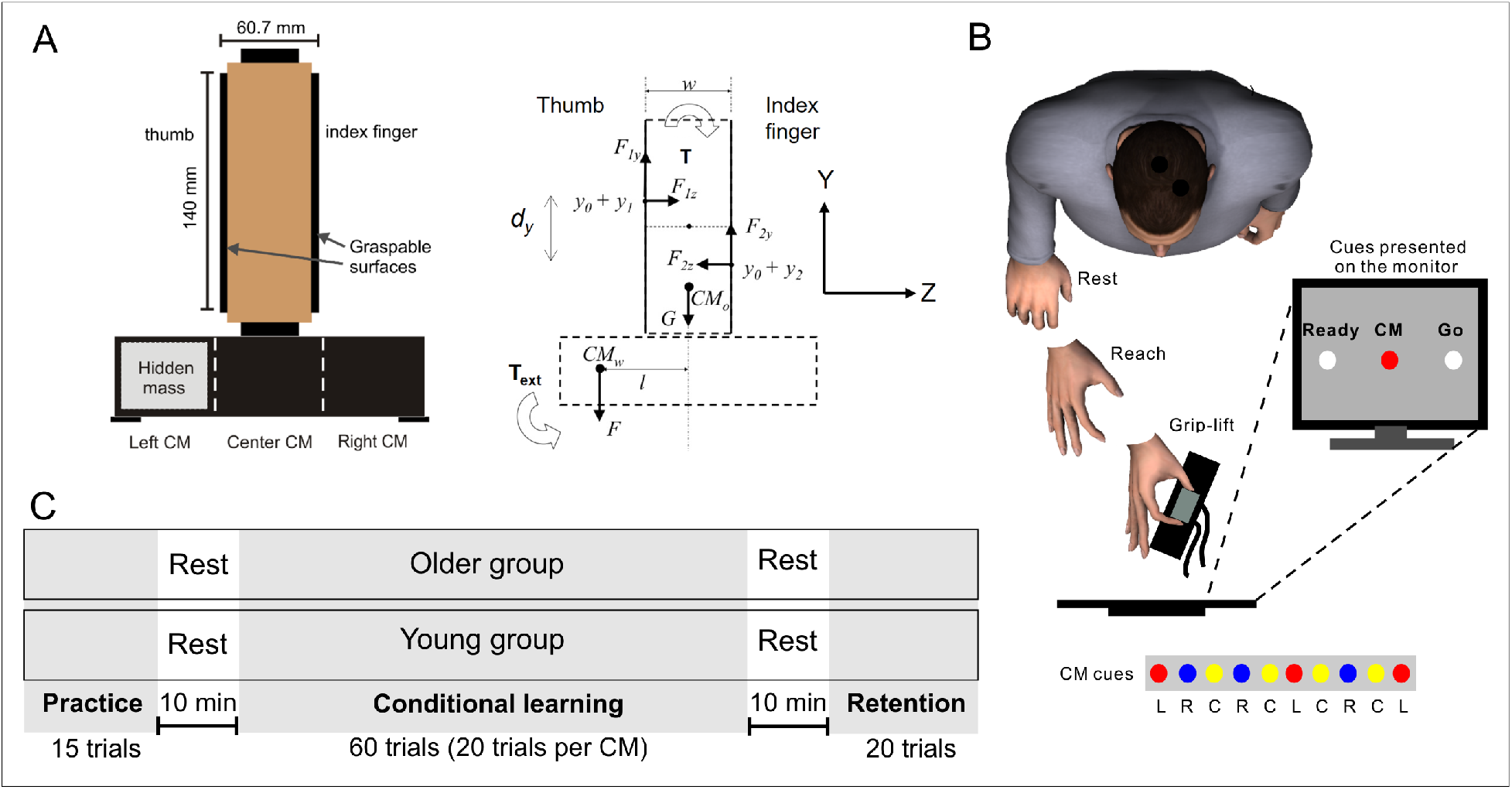
Experimental design. A: Grip device. B: Experimental setup. C: Experimental protocol.

### Experimental Procedure

#### Conditional visuomotor task

This task was adopted from our previous work (32). All experiments were conducted in a well-illuminated and quiet room. Subjects sat on a chair fitted with a platform designed to support their arm. The grip device was positioned 30 cm from the subject’s right hand on a table in front of the chair. Subjects were instructed to reach and grasp the grip device at a self-selected speed using their thumb and index fingertips of the right hand. They were instructed to lift the object to a height of ~10 cm while attempting to minimize object tilt, hold it for ~1 s, and replace it on the table. A computer monitor positioned in front of the subject was used to provide subjects with cues. For each trial, a series of three cues was presented to the subject (**Fig. 1B)**. First, a “ready” cue signaled the beginning of the trial. Next, a “CM” cue was presented at random delays (1-3 s) after the “ready” cue. For all but practice trials, this CM cue was a red, yellow, or blue in color and was arbitrarily associated with LCM, CCM, or RCM condition, respectively. For practice trials, the CM was always white in color and did not provide subjects with any information regarding the location of CM. Lastly, a “go” cue appeared on the monitor 1s after the CM cue to instruct subjects to begin the task. The task required subjects to associate a color cue with CM and using this association, learn to exert a torque in the appropriate direction and magnitude to counter the torque exerted by the cued CM.

### Experimental Protocol

#### Practice trials

Subject were familiarized with the experimental setup and the task by asking them to practice lifting the grip device and minimizing its roll. For the practice trials, the added mass (400 g) was placed in the right base compartment (RCM) of the grip device. Subjects were made aware that the added mass was going to be placed in the same compartment across all practice trials but the location of mass remained unknown to them. Subjects performed the task using a white cue which alerted the subjects about the upcoming task phase. Subjects performed 15 practice trials followed by a rest for ~10 min before proceeding with the conditional learning trials (**Fig. 1C)**.

#### Conditional learning trials

We altered the CM of the grip device on every trial by placing the mass in the right (RCM condition), center (CCM condition), or left compartment (LCM condition) at the base of the object (see **Fig. 1C**). The trials were pseudorandomized to avoid presenting the same CM over two consecutive trials. Subjects were informed about the trial-by-trial change in the CM location, but they were not explicitly told in which compartment the added mass would be across trials. Subjects were informed of the change in CM location using arbitrary color cues which were shown on a monitor directly in the line of sight of each subject. By trial and error, subjects were required to learn to associate the color cue with the CM location and select one of three possible responses: exertion of a clockwise, counterclockwise, or no torque, as per the color cue, and to lift the grip device while minimizing its tilt. Subjects received feedback about whether they successfully associated or failed to associate their response with the color cues (the stimulus) based on the direction and magnitude of the object tilt. Each CM condition was repeated 20 times (total 60 conditional learning trials). Before each trial, the experimenter changed the location of the added mass out of view of the subject. Since the practice trials were all completed with the RCM condition, and the sequence of the conditions was designed to avoid presenting the same CM on consecutive trials, the RCM condition was not presented on the first trial of the conditional learning block. Subjects were able to rest for 30 s between trials and this period included the time to change the location of the added mass. Following 60 conditional learning trials, subjects rested for ~10 min before beginning the retention trials.

#### Retention trials

Subjects were told to perform the conditional visuomotor task similar to the task performed in the conditional learning trials – by using the learned color-CM association. The location of the CM was, again, changed without the subjects’ knowledge by pseudorandom placement of the added mass in either of the three base compartments (right, center, and left) of the grip device. The condition presented in the last trial of the conditional learning block was not presented in the first trial of the retention block to avoid presentation of the same CM across two consecutive trials. Subjects rested between trials while the experimenters changed the location of the CM. The retention block consisted of 20 trials.

### Data Acquisition and Analysis

Conditional visuomotor learning of anticipatory control of dexterous manipulation was quantified using measurement and analysis of digit forces and positions at lift onset (32). Analysis of variables at object lift onset allows the quantification of subjects’ ability to coordinate digit forces and position based on recalling a force-position distribution associated with a specific color cue experienced in previous trials. After object lift onset, the object roll provides subjects with feedback about the extent to which the selected digit force and position were correctly associated with the color cue. The visual and haptic feedback about the magnitude and direction of object roll is used to drive the behavioral response on the next presentation of the same color cue. Across all practice trials, the CM location remained the same across consecutive trials (i.e. right center of mass) to allow subjects to use feedback of object roll to adjust digit force and placement on the following trial.

The following variables were analyzed: (1) Digit load force (LF) was defined as the vertical force component parallel to the grip surface produced by each digit to lift the object; (2) Normal force (F_N_) is the force component perpendicular to the grip surface; (3) Grip force (F_GF_), is the average of the normal forces produced by each digit; and (4) Digit center of pressure for each digit was defined as the vertical coordinate of the point of resultant force application by the digit on the grip surface. Using these variables, the following two variables were computed: (4) Torque and (5) Torque error (TE). In accordance with previous studies (31, 32), the torque generated by the subject was computed as:

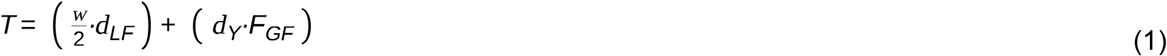

where d_LF_ is the difference between the LF of the thumb and index finger, d_Y_ is the vertical difference between the center of pressure of the thumb and index finger, F_GF_ is the grip force, w is the grip width, and 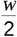 is the moment arm for the thumb and index finger LF (**Fig. 2A**). TE was defined as the absolute difference between the target torque (the torque required to counter the external torque) and the actual torque generated by the subject. The target torques for LCM and RCM conditions were 255 and −255 N*mm, respectively. (6) Peak object roll is the maximum roll occurring ~150 ms after object lift-off.

**Figure 2.**
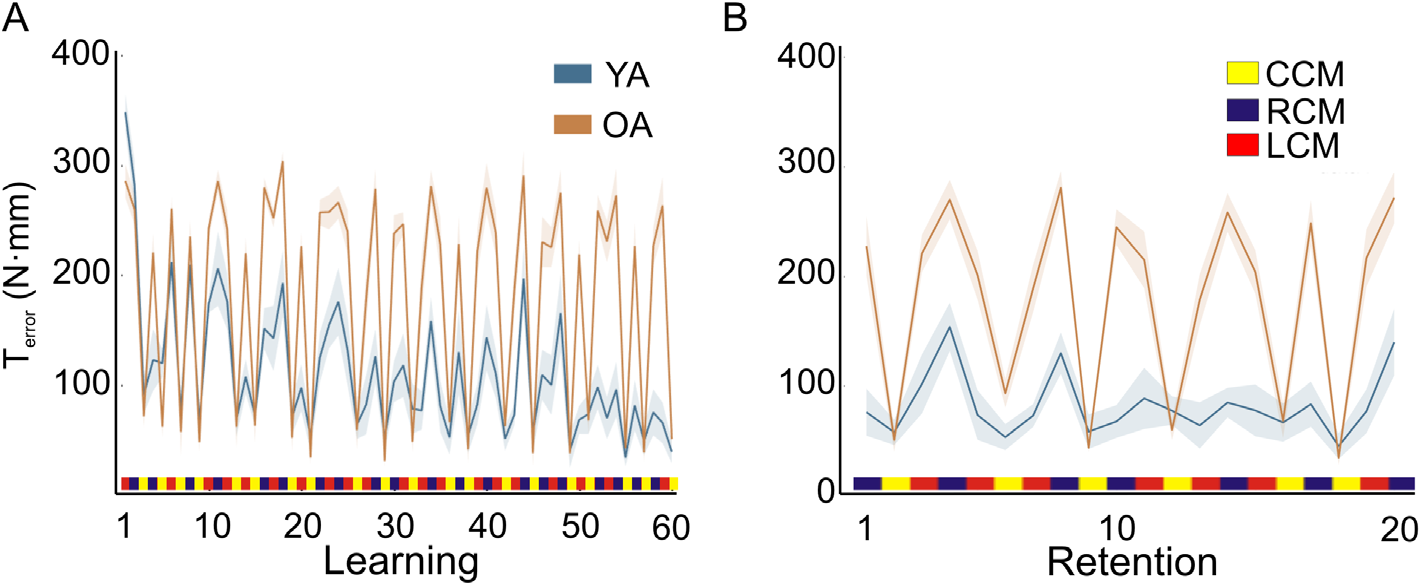
Conditional learning and retention. Torque error (TE) plotted as a function of trial, during A: conditional learning and B: retention among older adults (OA) and young adults (YA). The CMs presented in a pseudorandom order across trials are denoted by different colors on the horizontal axis. Data are averages (± SEM) of all subjects for both plots.

All of the above variables, except peak roll, were computed at the time of object lift onset to quantify anticipatory control of manipulation (31, 32). Object lift onset was defined as the time at which the vertical position of the grip device crossed and remained above a threshold (mean + 2SD of the baseline) for 200 ms (44).

### Statistical Analyses

For the practice block, we measured each subjects’ ability to apply a torque to help minimize the roll of the grip device by comparing the first practice trial with the average of the last five practice trials. For the conditional learning block, we averaged the TE and the object roll measurements across three successive learning trials (average of trials 1-3, average of trials 4-6, average of trials 7-9, etc.). For the retention block, we averaged the TE and the object roll measurements across three successive trials for the first 18 trials. We performed repeated measures analysis of variance (ANOVA; ***α*** = 0.05) on TE, roll, d_Y_, d_LF_, and F_GF_ with GROUP (Older, Young) as a between-subject factor and one or more of the following within subject factors: *Bin*_*learning*_ (20 levels, bin_learning_ 1 to 20), *Bin*_*retention*_ (6 levels, bin_retention_ 1 to 6), *CM* (LCM, RCM, CCM), *Trial* (1–20), and *Practice Trial* (first, average of the last 5 trials). For the conditional learning block, we performed a post hoc between-group comparisons for bin_learning_ 1 and bin_learning_ 20, which represent the first and last exposure to each CM condition, respectively. We applied Huynh-Feldt corrections when sphericity assumption was violated. Post hoc comparisons using paired t-tests were performed with Bonferroni corrections, if needed. ANOVA and posthoc analyses were performed using SPSS software version 22.0 (IBM, USA).

We also report estimation statistics that describe the magnitude and precision of the effect size (6, 8). A total of 5000 bootstrap samples were taken (with replacement), and the resampling distribution of difference in mean (i.e. effect size) was determined (20). A bias-corrected bootstrap 95% confidence interval (CI) was constructed from the resampling distribution. The data was analyzed and figures were produced using the estimation coding software (20). The p-value obtained from the two-sided permutation t-test (***α*** = 0.05) provided the likelihood of observing the effect size, if the null hypothesis of zero difference is true (**Table 1**).

**Table 1.**
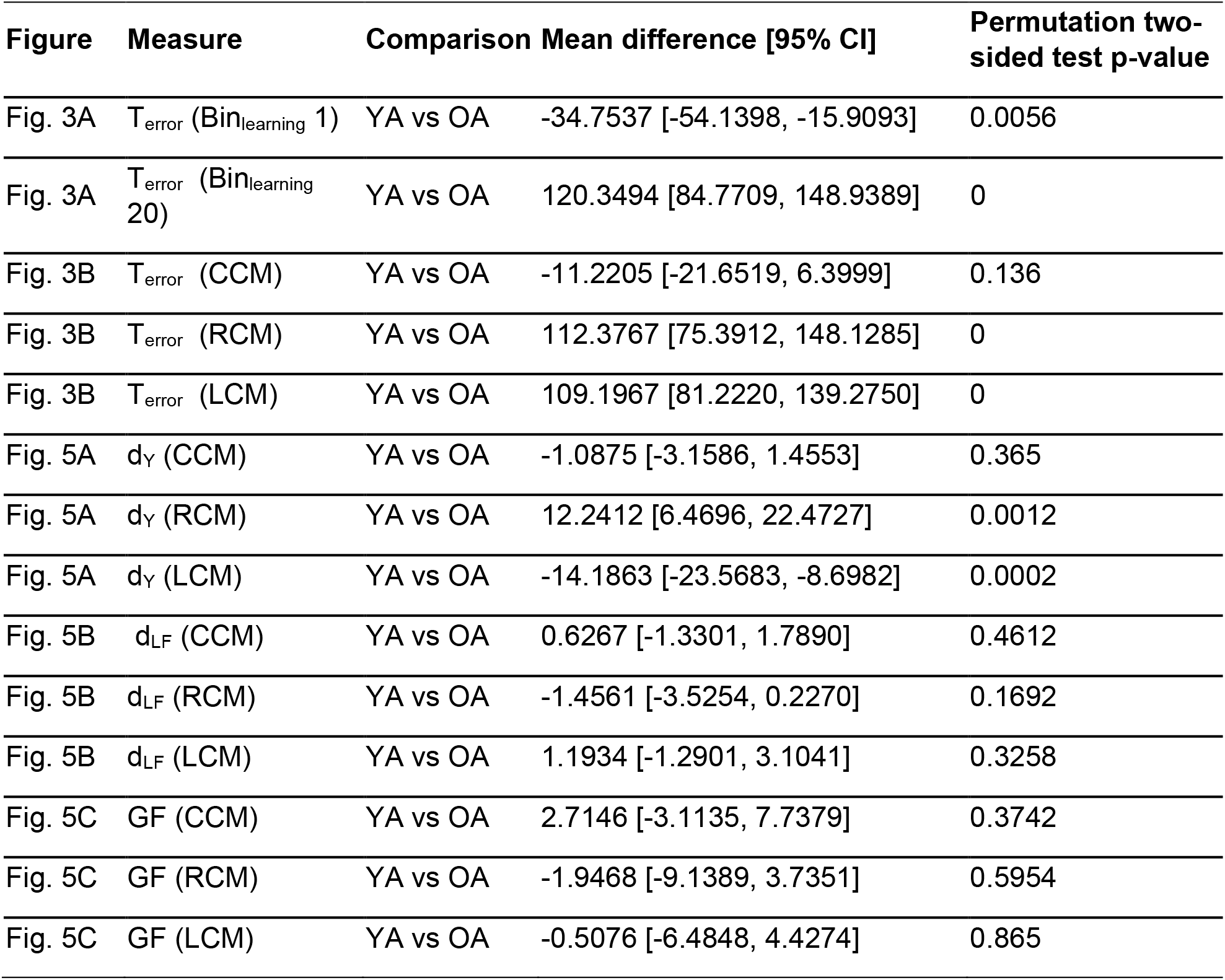
Estimation statistics organized by figure panel. The type of measurement, comparison, mean difference with 95% confidence interval (CI), and the p-value for the permutation two-sided test are provided.

| Figure | Measure | Comparison | Mean difference [95% CI] | Permutation two-sided test p-value |
| --- | --- | --- | --- | --- |
| Fig. 3A | $T_{\text{error}}$ ($\text{Bin}_{\text{learning}} 1$) | YA vs OA | -34.7537 [-54.1398, -15.9093] | 0.0056 |
| Fig. 3A | $T_{\text{error}}$ ($\text{Bin}_{\text{learning}} 20$) | YA vs OA | 120.3494 [84.7709, 148.9389] | 0 |
| Fig. 3B | $T_{\text{error}}$ (CCM) | YA vs OA | -11.2205 [-21.6519, 6.3999] | 0.136 |
| Fig. 3B | $T_{\text{error}}$ (RCM) | YA vs OA | 112.3767 [75.3912, 148.1285] | 0 |
| Fig. 3B | $T_{\text{error}}$ (LCM) | YA vs OA | 109.1967 [81.2220, 139.2750] | 0 |
| Fig. 5A | $d_Y$ (CCM) | YA vs OA | -1.0875 [-3.1586, 1.4553] | 0.365 |
| Fig. 5A | $d_Y$ (RCM) | YA vs OA | 12.2412 [6.4696, 22.4727] | 0.0012 |
| Fig. 5A | $d_Y$ (LCM) | YA vs OA | -14.1863 [-23.5683, -8.6982] | 0.0002 |
| Fig. 5B | $d_{\text{LF}}$ (CCM) | YA vs OA | 0.6267 [-1.3301, 1.7890] | 0.4612 |
| Fig. 5B | $d_{\text{LF}}$ (RCM) | YA vs OA | -1.4561 [-3.5254, 0.2270] | 0.1692 |
| Fig. 5B | $d_{\text{LF}}$ (LCM) | YA vs OA | 1.1934 [-1.2901, 3.1041] | 0.3258 |
| Fig. 5C | GF (CCM) | YA vs OA | 2.7146 [-3.1135, 7.7379] | 0.3742 |
| Fig. 5C | GF (RCM) | YA vs OA | -1.9468 [-9.1389, 3.7351] | 0.5954 |
| Fig. 5C | GF (LCM) | YA vs OA | -0.5076 [-6.4848, 4.4274] | 0.865 |

## RESULTS

### Practice

Both older and young adults practiced the conditional visuomotor task to familiarize with the task requirement of object roll minimization (RCM only). Subjects were required to produce a torque on the object prior to lift onset that had to be equal in magnitude but opposite in direction to the external torque generated by the added mass (target torque = −255 N·mm) to minimize object roll. That is, subjects were required to anticipate, rather than react to, the external torque, through consecutive lifts. As they were unaware of the CM location on the first practice trial and because the object is visually symmetrical, both older and young adults exerted no or negligible torque on the object. This led to large torque error (TE - Young adults: 210 ±30.11 N·mm; Older adults: 276.5 ±15.6 N·mm) and object roll (Young adults: 12.18° ±2.9°; Older adults: 11.36° ±1.34°). Importantly, subjects were able to reduce TE and object roll over the remaining practice trials (TE - Young adults: 44.95 ±5.26 N·mm; Older adults: 118 ±10.2 N·mm and object roll - Young adults: 3.67° ±0.29°; Older adults: 3.12° ±0.33°). Both older adults and young adults were able to significantly reduce TE in a similar manner through practice (main effect of Practice Trial: F_1, 18_ = 78.68; p < 0.001, η_p_^2^ = 0.81; no significant Group × Practice Trial interaction: p = 0.87). However, across practice trials, older adults demonstrated significantly greater TE than young adults (main effect of Group: F_1, 18_ = 15.75; p = 0.001, η_p_^2^ = 0.47). Similar overall results were found for object roll (main effect of Practice Trial: F_1, 18_ = 28.84, p < 0.0001, η_p_^2^ = 0.62; no significant Group × Practice Trial interaction: p = 0.93; no main effect of Group: p = 0.69).

These findings, along with previous studies (23, 24, 31, 32), demonstrated that Tcom is a valid predictor of manipulation performance, i.e., object roll. Specifically, as subjects learn the appropriate Tcom required to minimize object roll, peak object roll negatively correlates with the magnitude of Tcom. We also note that the results of the analysis of peak object roll and Tcom were identical across young and older groups. Because both variables capture two interrelated phenomena associated with learning dexterous manipulation, for the sake of brevity we will use only TE as the measure of anticipatory grasp control throughout the rest of the manuscript.

Both young and older adults modulated their digit position over the practice trials (main effect of Practice Trial: F_1, 18_ = 5.99; p = 0.025, η_p_^2^ = 0.25; no significant Group × Practice Trial interaction: p = 0.48; no main effect of Group: p = 0.59). On the first practice trial, all subjects placed their thumb and index finger at nearly same location on their corresponding graspable surfaces (Young adults: −1.41 ±1.41 mm and Older adults: −1.44 ±1.56 mm). All subjects learned to position their index fingertip higher than the thumb with repeated exposure to the object with an external mass on the right side (Average of last five trials - Young adults: −7.3 ±3.75 mm and Older adults: −4.7 ±1.1 mm). Similarly, both young and older adults learned to apply significantly higher load force with their index fingertip than the thumb over the practice trials (main effect of Practice Trial: F_1, 18_ = 37.93; p < 0.001, η_p_^2^ = 0.68; no significant Group × Practice Trial interaction: p = 0.47; no main effect of Group: p = 0.07). On the first trial, both groups exerted similar load force with their digits (Young adults: −0.76 ±0.87 N and Older adults: −0.75 ±0.6 N). During the later practice trials, both groups applied higher load force with their index finger than the thumb (Young adults: −3.9 ±0.9 N and Older adults: −1.77 ±1.69 N). Both groups exerted greater grip force on the object with repeated practice (main effect of Practice Trial: F_1, 18_ = 6.2; p = 0.023, η_p_^2^ = 0.26; no significant Group × Practice Trial interaction: p = 0.48; no main effect of Group: p = 0.12). Young adults increased their grip force from 11.01 ±1.65 N to 14.07 ±1.62 N while older adults increased their grip force from 12.92 ±2.18 N to 18.47 ±1.83 N across the practice trials. These results indicate that both the groups showed intact ability to modulate the components of TE i.e. digit position, load force, and grip force during practice trials. It is likely that higher TE among older adults could underlie negligible, non-significant changes in load force and grip force and their associated multiplicative effects on the application of torque (see eq. 1).

### Conditional visuomotor learning in older adults

Both young and older adults were instructed to perform a conditional visuomotor task using the object whose CM was altered on a trial-by-trial basis in a pseudorandom fashion, thus resulting in three CM conditions: RCM, CCM, and LCM (**Fig. 1**). They were also informed about the change in CM of the object through arbitrary color cues where red, blue and yellow colors corresponded to LCM, RCM, and CCM conditions respectively. However, they had no *apriori* information about the color – object CM association. By trial and error, all adults were required to learn to associate the color cue with object CM and select accurate torque, according to the identified cue, and to lift the object while minimizing object tilt. Minimization of object tilt required application of torque on the object that approached the target external torque. The direction and magnitude of object tilt provided feedback to subjects about success or failure of their stimulus-response association. The trial sequence was designed to inform individuals about color-object CM association in the first three trials. As both young and older adults were unaware of the association between the color and object CM during the first two trials (LCM followed by RCM), they produced very large TE during these trials. On trial 3, we observed a sudden reduction in TE as compared to trial 1. Note that the same CM condition was not repeated over two consecutive trials. Therefore, presentation of the third color cue on trial 3 would have allowed subjects to infer, by elimination, the upcoming CM condition. However, an alternative explanation for the drop in TE on trial 3 is that subjects did not attempt to infer object CM in the initial exposure to the color cues, but rather that they exerted, as a default, little to no torque in the early trials. Furthermore, although subjects were exposed to all color-object CM associations by trial 3, they continued to produce a fairly large TE. Young adults gradually reduced TE over 60 learning trials. In contrast, older adults failed to reduce TE over 60 learning trials.

To quantify the time course of conditional learning, we focused our analysis on the trial ‘bins’ (average of 3 trials; (32)). We found that the reduction in TE across the conditional learning trials was significantly smaller in older adults when compared with young adults (significant Group × Bin_learning_ interaction: F_19, 342_ = 9.74, p < 0.0001, η_p_^2^ = 0.35; **Fig. 2A**). Specifically, at the end of learning trials (i.e. bin_learning_ 20), we found that older adults showed significantly greater TE than young adults (t_18_ = 6.98; p < 0.0001; **Fig. 3A**). For bin_learning_ 1, older adults showed significantly smaller TE than young adults (t_18_ = 3.375; p = 0.003). We also found a significant main effect of Group (F_1, 18_ = 39.56, p < 0.0001, η_p_^2^ = 0.69) and Bin (F = 37.76, p < 0.0001, η_p_^2^ = 0.68). An exponential fit of the TE data averaged across subjects (y = a·e^bx^ + c) resulted in a half-life of 45 trials and 252 trials in young and older adults, respectively (significant difference in exponential coefficients; t_18_ = 4.22; p = 0.001). These findings suggest that older adults were unable to learn to accurately produce the torque using color cues about CM location to the same extent as young adults by the 60^th^ conditional learning trial. Following the conditional learning trials, young adults were able to recall the association between arbitrary color cues and object CM to select the torque appropriate to a given CM learned during the learning trials (see **Fig. 2B**).

**Figure 3.**
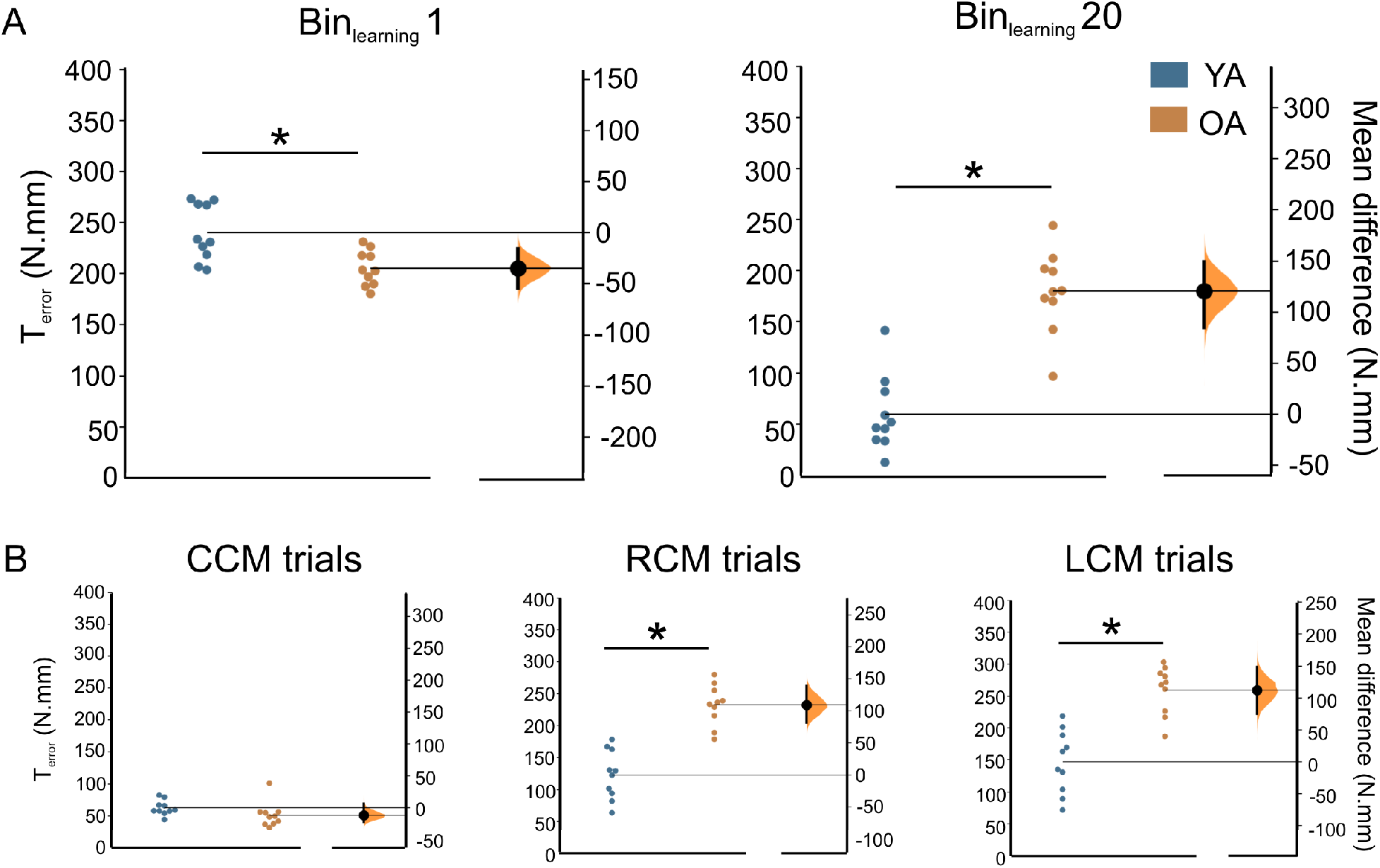
Estimation plots for torque error (TE). A: TE for bin_learning_ 1 (left) and bin_learing_ 20 (right) during conditional learning trials in older adults (OA) and young adults (YA). B: TE is shown for each object CM for older adults (OA) and young adults (YA). Data for individual subjects (solid circles) in both groups are plotted on the left axes and the mean difference between groups is plotted on the right axes as a bootstrap resampling distribution. The mean difference is depicted as a large black dot with the 95% confidence interval indicated by the ends of the vertical error bar. Asterisks denote significant differences (p < 0.05).

We expected the effect of aging on the production of torque (close to the target torque) during conditional learning to also vary across CM conditions. We found that the reduction in TE for older and young adults differed across CM conditions (significant Group × CM interaction: F_2, 36_ = 31.88, p < 0.0001, η_p_^2^ = 0.64; **Fig. 3B**). Furthermore, we found that older adults demonstrated greater TE than young adults for the RCM (t_18_ = 5.73; p < 0.0001) and the LCM (t_18_ = 6.94; p < 0.0001) conditions, but not for the CCM condition (t_18_ = 1.58; p = 0.14).

Next, we present findings from the individual behavioral variables that constituted the torque exerted on the object (see equation 1) to further understand the effects of aging on conditional visuomotor learning.

### Modulation of digit position during conditional learning

Older adults failed to use the correct modulation of index finger and thumb based on the arbitrary color cues about object CM when compared with young adults (Group × CM interaction: F_1.09., 19.62_ = 12.89, p = 0.002; η_p_^2^ = 0.42; but no main effect of Group: F_1, 18_ = 0.57, p = 0.46; **Figs. 4A** **and** **5A**). That is, young adults but not older adults modulated the relative placement of the thumb and index finger on the object (d_Y_) to the CM location during conditional learning. However, the modulation in d_Y_ across learning trials was mainly observed for the RCM and LCM conditions (significance CM × Trial interaction: F_6.04, 108.79_ = 7.8; p < 0.0001; main effect of Trials: F_19, 342_ = 5.04, p < 0.0001; η_p_^2^ = 0.22). For the CCM condition, both young and older adults placed their thumb and index finger at nearly same location on their corresponding graspable surfaces (t_18_ = 0.89; p = 0.39; **Figs. 4A** **and** **5A**). Importantly, for the RCM condition across the conditional learning trials, young adults chose to place their index finger higher than thumb on the object while older adults did not (t_18_ = −3.02; p = 0.007; **Figs. 4A** **and** **5A**). For the LCM condition, young adults chose to place their thumb higher than index finger on the object while older adults did not (t_18_ = 3.72; p = 0.002; **Figs. 4A** **and** **5A**). The observed modulation of d_Y_ in young adults provide objective evidence of their ability to identify the CM of the object using color cues and modulate digit placement accordingly. Such CM location-dependent modulation of digit placement was absent in older adults.

**Figure 4.**
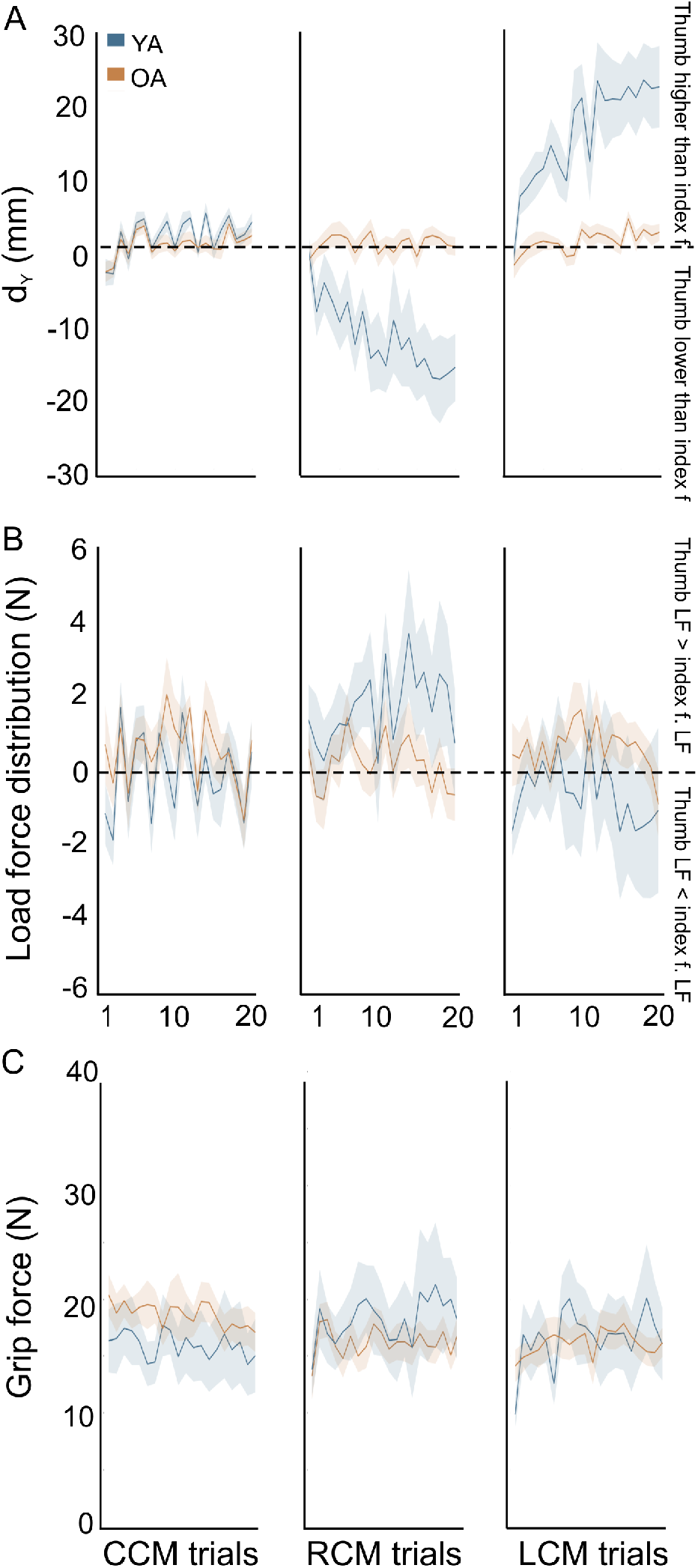
Digit placement, load force distribution, and grip force during conditional learning. *A:* Vertical distance between thumb and index finger center of pressure (d_Y_) is plotted as a function of trial for each object CM for older adults (OA) and young adults (YA) *B:* Difference between thumb and index finger load force (d_LF_) is plotted as a function of trial for each object CM for older adults (OA) and young adults (YA). *C:* Grip force (GF) is plotted as a function of trial for each object CM for older adults (OA) and young adults (YA). Data are averages of all subjects (± SEM).

**Figure 5.**
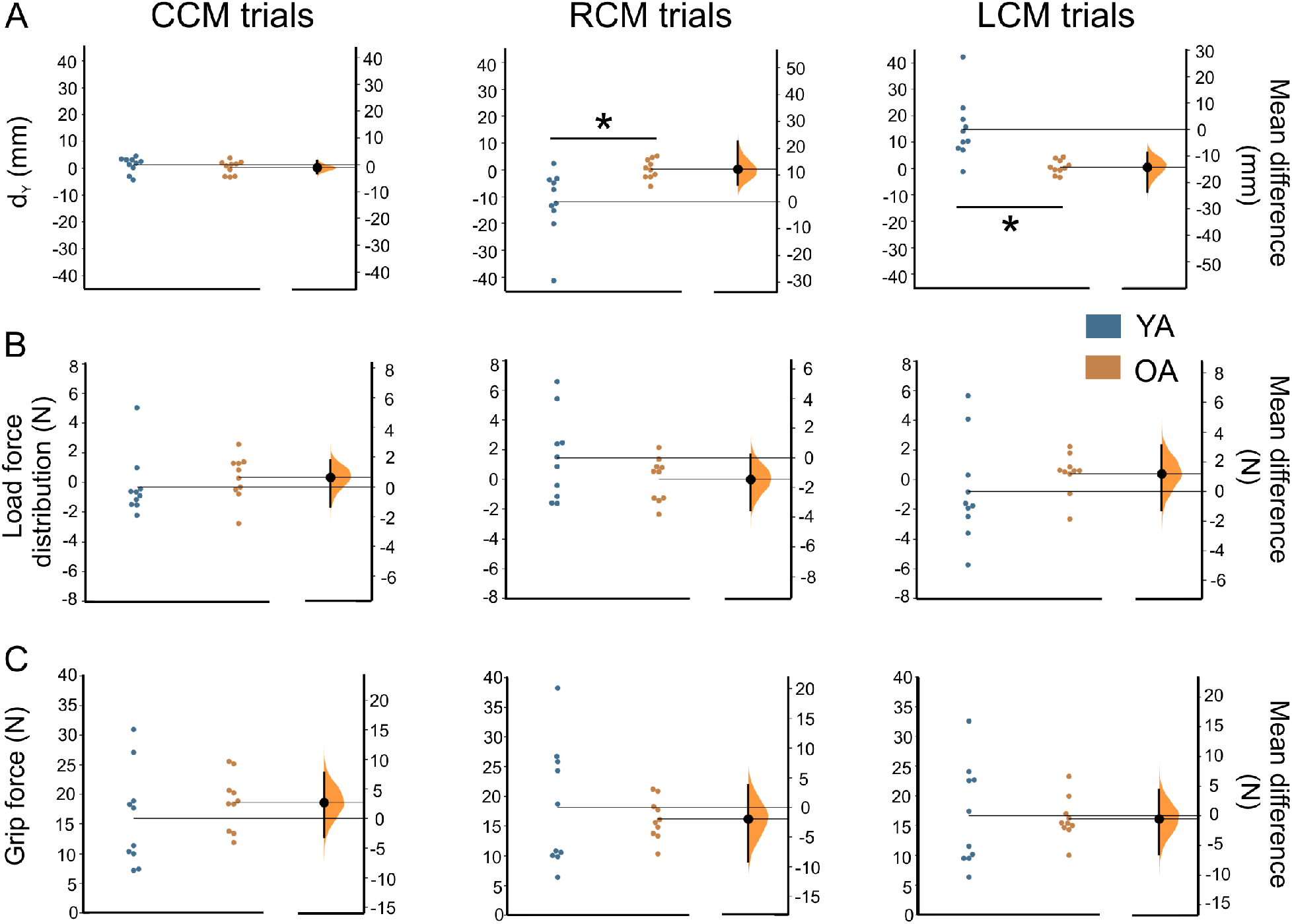
Estimation plots for digit placement, load force distribution, and grip force during conditional learning. *A:* Vertical distance between thumb and index finger center of pressure (d_Y_) is shown for each object CM for older adults (OA) and young adults (YA). *B:* Difference between thumb and index finger load force (d_LF_) is shown for each object CM for older adults (OA) and young adults (YA). *C:* Grip force (GF) is shown for each object CM for older adults (OA) and young adults (YA). Data for individual subjects (solid circles) in both groups are plotted on the left axes and the mean difference between groups is plotted on the right axes as a bootstrap resampling distribution. The mean difference is depicted as a large black dot with the 95% confidence interval indicated by the ends of the vertical error bar. Asterisks denote significant differences (p ≤ 0.05).

### Modulation of digit forces during conditional learning

Modulation of d_LF_ to CM location was significantly different in older adults when compared with young adults (Group × CM interaction: F_2, 36_ = 3.38; p = 0.04; η_p_^2^ = 0.16; but no main effect of Group: F_1, 18_ = 0.024; p = 0.88; **Figs. 4B** **and** **5B**). However, we failed to observe significant differences in d_LF_ between young and older adults for RCM (t_18_ = 1.436; p = 0.17), LCM (t_18_ = −1.03; p = 0.32), and CCM (t_18_ = −0.78; p = 0.45). The magnitude of d_LF_ changed across trials (main effect of Trials: F_19, 342_ = 2.8; p < 0.0001; η_p_^2^ = 0.13). However, the change in d_LF_ across trials was similar across CM conditions (no CM × Trial interaction: F_38, 684_ = 1.33; p = 0.09).

Similarly, although modulation of grip force (F_GF_) to CM location was significantly different in older adults versus young adults (Group × CM interaction: F_2, 36_ = 10.21; p < 0.0001; η_p_^2^ = 0.36; but no main effect of Group: F_1, 18_ = 0.001; p = 0.98), posthoc comparisons failed to observe significant differences between two groups (**Figs. 4C** **and** **5C**, RCM: t_18_ = 0.57; p = 0.58, LCM: t_18_ = 0.17; p = 0.86, and CCM: t_18_ = −0.91; p = 0.37). Grip force did not significantly change across trials (significant CM × Trial interaction: F_38, 684_ = 1.91, p = 0.001, η_p_^2^ = 0.096, but no main effect of Trial: F_19, 342_ = 0.99; p = 0.47).

Table 1 on estimation statistics reports the group mean difference with corresponding 95% CI (effect size), and the p-value for the permutation two-sided test. The findings of the estimation statistics were found to be consistent with the ANOVA findings.

## DISCUSSION

We investigated whether older adults are able to modulate digit position based on arbitrary visual cues about object center of mass (CM) for dexterous manipulation. A novel finding of this study is that the older adults failed to learn to use arbitrary visual cues about object CM for anticipatory control of digit position, when compared with young adults. Consistent with (7), older adults did not learn to scale their digit forces based on arbitrary color cues about object CM location. A lack of modulation of digit position and digit forces in older adults was reflected in their impaired ability to minimize the error in compensatory torque exerted on the object across the conditional learning trials. Thus, we observed a slower rate of conditional visuomotor learning in older adults when compared with young adults. Importantly, this impairment was specific to conditional learning as the older adults showed an intact ability to learn to modulate digit position as well as digit force based on implicit knowledge of object property. We discuss these findings in the context of age-related deficits in processes essential for conditional visuomotor learning and bridge this understanding with previously reported neurophysiological mechanisms underlying dexterous manipulation.

### Attentional Mechanisms, Aging, and Conditional Visuomotor Learning

The conditional learning paradigm necessitates formation of association between the stimulus and its predictive relevance in a given task (26, 32). This process is argued to rely on the modulation of attentional mechanisms to extract the predictive information from the relevant stimulus while ignoring irrelevant or distracting stimuli (26, 40). Consequently, an impairment in the ability to modulate attention could impair conditional learning. In our study, however, it is less likely that older adults were impaired in attending to the visual color cues about object CM. The experimenters ensured subjects’ compliance with instructional cues during the sessions, e.g. initiating a reach only after the presentation of *go* cue. The practice trials gave subjects an opportunity to familiarize themselves with the visual cues, the experimental setup and the laboratory ambience. Both young and older adults were able to follow the instructional cues (*ready, task, go)* to successfully perform the object manipulation task during practice trials. After every associative visuomotor learning trial, subjects were made aware of the change in the location of external mass because these changes were done in their presence, although hidden from their view. Importantly, subjects accurately reported the change in color of the *task* cue after every learning trial. Moreover, a recent study systematically investigated whether an impairment in attentional network with aging could affect conditional learning by instructing healthy older and younger adults to perform category learning and dot-probe tasks (26). Despite lower accuracy in predicting association pairs on the category learning task, older adults learned to modulate their attention toward predictive stimuli while directing it away from the irrelevant ones, similar to the young adults. The findings from the dot-probe task provided a more direct evidence during which the older adults rapidly responded to the dot probe when cued by a predictive stimulus versus a non-predictive stimulus. Thus, the ability to modulate attentional processes to derive predictive information from the arbitrary cues was found to be intact in older adults (26). Our study involved a similar process of associating a color cue-based stimulus to the object CM location relevant to accomplish the task requirement. It is, therefore, likely that age-related impairments in the anticipatory modulation of digit position and forces found in this study were due not to older adults’ inability to modulate the attentional mechanisms during conditional learning.

### Connecting the dots: Effects of Aging on Conditional Visuomotor Learning during Dexterous manipulation

We found that older adults were slower in learning novel color-torque associations to lift the object while minimizing object tilt (significant *Group* × *Conditional learning bins* interaction effect). This is in agreement with studies demonstrating impaired learning using explicit cues in older adults (7, 11, 37, 39, 41, 45). Notably, the observed impairment in the application of compensatory torque on the object in older adults during conditional learning trials was not due to poor motor execution as these adults, similar to young adults, successfully learned to apply accurate compensatory torque on the object during practice trials (significant learning effect but no *Group* × *Practice trials* interaction effect). Therefore, the ability to acquire representations of the object’s properties through arbitrary visuomotor associations is likely impaired in old age (7). We consider possible mechanisms underlying the age-dependent impairment in conditional learning during dexterous manipulation.

During conditional learning trials, subjects had to select one of three possible responses (i.e., exertion of clockwise, counterclockwise, or no torque) to lift the object while minimizing its tilt across 60 trials. Selection of a response depends on a visual identification of the color cue that provides knowledge about the object’s CM location (2, 32). It is possible that this visual identification of object CM location was impaired, which might have affected the selection of accurate compensatory torque. Alternatively, the identification of visual cue was intact but the subsequent transformation of this information into motor commands was impaired in older adults (30). Older adults exerted erroneous compensatory torque on the object across the conditional learning trials. The erroneous compensatory torque was primarily observed for the RCM and LCM trials and not for the CCM trials. The application of accurate torque, a requirement during the RCM and LCM trials, is a function of correct spatial distribution of digit contact points (d_Y_) and CM-appropriate distribution of load forces across thumb and index finger (d_LF_) on the object (12, 13, 32, 44). Young adults were found to place the index fingertip higher than the thumb when cued about a right CM, and lower than the thumb when cued about a left CM as conditional learning progressed, a finding consistent with (32). This result is important as it indicates that young adults could correctly identify object CM, an ability that would have allowed them to exert a torque with the appropriate direction on the object. In contrast, older adults failed to show modulation of digit position using arbitrary visual cues about object CM location throughout the conditional learning trials (**Figs. 4** **and** **5**). That is, older adults placed their thumb and index finger at nearly same location on their corresponding graspable surfaces irrespective of object CM and did not demonstrate a tendency to learn the anticipatory modulation of digit position using color cues over the learning trials. This is a novel and important finding as the same group of older adults were able to learn to successfully and anticipatorily modulate their digit contact points based on implicit knowledge of object property during practice trials. For all practice trials, the external mass remained in the right compartment of the base of object. Thus, the CM condition during practice trials was analogous to the CM condition (i.e. RCM) during conditional learning trials, except that the latter condition introduced the color-coded cue associations which were interspersed with the other two object CM conditions (LCM and CCM). Importantly, the finding from the practice trials are consistent with our previous report (30) of intact ability of older adults to modulate grasp parameters based on somatosensory and visual information about task performance. It should be noted that practice with right-sided mass did not provide an undue advantage during conditional learning trials because the deficits in digit position modulation were similar across RCM and LCM conditions (**Figs. 4** **and** **5**).

In addition to deficits in anticipatory modulation of digit position, older adults were impaired in scaling their digit forces based on color cues about object CM when compared with young adults. Cole and Rotella (7) have reported that older adults are impaired in acquiring the ability to select digit forces specific to object property based on arbitrary color cues about object weight and texture at contact surfaces. In addition to confirming this finding, our results point toward a deficit in an important feature of dexterous manipulation - modulation of digit forces as a function of digit position - in older adults. In young adults, digit force modulation (e.g. more positive d_LF_ for later RCM trials and more negative for later LCM trials; see **Fig. 4B**) was found to compensate for large modulation of digit position (e.g. more negative d_Y_ for later RCM trials and more positive for later LCM trials). The unusually large modulation of digit position, if left uncorrected, would have resulted in larger than necessary compensatory torque application, thus tilting the object in the opposite direction. In contrast, older adults failed to modulate digit forces to account for the lack of anticipatory control of digit position during conditional learning trials.

Inability to modulate digit position and forces using the color cues about object CM and the lack of digit force-to-position compensation during conditional learning trials led to erroneous application of compensatory torque on the object. Our previous study (31) used a grasp device and torque requirement (viz. the RCM condition) similar to the one referred in the current study, but with an aim to identify neural centers processing different grasp parameters during dexterous manipulation using implicit knowledge of the object CM location. We found that the role of primary somatosensory cortex (S1) and M1 was critical to modulate digit position and forces for accurate application of compensatory torque to minimize the object tilt. Additionally, Zach and colleagues (43) have shown that M1 neurons in non-human primates develop novel representations of behaviorally relevant features such as arbitrary color cues during conditional learning. It is likely that the age-dependent altered activation within M1 and S1, as reported by several studies (3, 14, 18, 19, 21), could underlie the impaired processing of digit position and forces thereby, affecting response selection during the conditional learning block.

The selected response during conditional learning needs to be monitored so that feedback about the behavioral consequences of the current response could inform subsequent responses for similar stimulus presentations – either by modifying it on future stimulus presentations if a selection error was made, or repeating it if the selected response was correct (7, 32). Object tilt occurring after lift onset provides subjects with feedback about the extent to which the selected force-position distribution was correctly associated with the color cue. Visual and haptic feedback about the magnitude and direction of object roll is then used to drive the response on the next presentation of the same color cue. In case of practice trials where the CM condition (i.e. RCM) remained the same across consecutive trials, older and young adults alike could use feedback of object tilt to control digit placement and force distribution on the subsequent trial. This finding suggests that older adults had an intact ability to monitor behavioral consequences of the response during learning trials, a finding consistent with (30). However, the CM conditions were pseudorandomized during conditional learning trials, which required the subjects to remember this information and use it during next presentation of the same color cue. It is possible that older adults were unable to store this information in memory and/or retrieve this information to guide behavior on subsequent stimulus presentations. The hippocampal connection with prefrontal cortex is found to be critical for aiding the formation of novel associations during a conditional learning task (4). Neurophysiological evidence based on non-human primates suggests that aging accompanies reduction in working memory capacity due to reduced hippocampal activity over the lifespan (2, 7, 11). An age-related reduction in bold-related activity in prefrontal regions has been found during the formation of sensorimotor mappings (10) Additionally, our previous study (32) using a similar conditional visuomotor paradigm showed that contralateral dorsal premotor cortex (PMd) plays an important role during the formation of novel associations in young adults. Mainly, disruption of PMd using continuous theta burst transcranial magnetic stimulation lead to a slower rate of learning, but did not affect the recall of a learned conditional visuomotor behaviour. The age-related decrement in the acquisition of novel associations might be linked to age-related changes in hippocampus, basal ganglia, prefrontal, premotor and motor regions (10, 15, 29, 29, 33, 38, 42). Future studies investigating a broader constellation of factors (17) influencing the age-dependent modulation in memory mechanisms hold promise to elucidate the holistic impact of aging on conditional visuomotor learning.

Conditional learning is believed to be complete when the error in motor response selection cued by an arbitrary stimulus is reduced and reaches a plateau (7, 32). Using this learned stimulus-response association, subjects are able to select an effective motor response when a familiar stimulus is presented (see **Fig. 2B**). Although we are not able to comment on older adults’ ability to recall as the conditional visuomotor task was not learned, there is evidence for absence of deficit in recalling a learned behavior. Fisk and Warr (11) have shown that the older adults were able to accurately recall previously learned associations despite being impaired in acquiring new associations between individual cursor movements and the corresponding keys over a number of learning trials as indicated by the greater number of incorrect attempts when compared with young adults. Similarly, older adults were found to be impaired in acquiring novel stimulus-response association and not in the retention of learned associations (39).

Overall, our findings suggest that the older adults’ inability to predictively modulate digit position and digit force based on arbitrary color cues about object property might be due to age-related alterations in the conditional learning processes such as stimulus identification, response selection, and/or memory formation. These processing decrements might be due to age-related changes in the cortical-subcortical network involving prefrontal, premotor, sensorimotor, hippocampus, and basal ganglia regions.

## ACKNOWLEDGMENTS

This work was supported by the University of Houston, Division of Research High Priority Area Research Seed Grant to PJP. We thank Kevin Nguyen for assistance with data collection.

## Notes

### Competing Interest Statement

The authors have declared no competing interest.

## REFERENCES

1. Ameli M, Kemper F, Sarfeld AS, Kessler J, Fink GR, Nowak DA. Arbitrary visuo-motor mapping during object manipulation in mild cognitive impairment and Alzheimer’s disease: A pilot study. Clin Neurol Neurosurg 113: 453–458, 2011.

2. Asaad WF, Rainer G, Miller EK. Neural activity in the primate prefrontal cortex during associative learning. Neuron 21: 1399–407, 1998.

3. Bernard JA, Seidler RD. Evidence for motor cortex dedifferentiation in older adults. Neurobiol Aging 33: 1890–1899, 2012.

4. Brincat SL, Miller EK. Frequency-specific hippocampal-prefrontal interactions during associative learning. Nat Neurosci 18: 576–581, 2015.

5. Chouinard PA, Leonard G, Paus T. Role of the primary motor and dorsal premotor cortices in the anticipation of forces during object lifting. J Neurosci 25: 2277–84, 2005.

6. Cohen J. The earth is round (p <. 05). What If There Were No Significance Tests? Class. Ed. (2016). doi: 10.4324/9781315629049-14.

7. Cole KJ, Rotella DL. Old age impairs the use of arbitrary visual cues for predictive control of fingertip forces during grasp. Exp Brain Res 143: 35–41, 2002.

8. Cumming G, Calin-Jageman RJ. Introduction to the New Statistics: Estimation, Open Science, and Beyond. Routledge, 2016.

9. Davare M, Parikh PJ, Santello M. Sensorimotor uncertainty modulates corticospinal excitability during skilled object manipulation. J Neurophysiol 121: 1162–1170, 2019.

10. Dennis NA, Hayes SM, Prince SE, Madden DJ, A S, Cabeza R. Effects of Aging on the Neural Correlates of Successful Item and Source Memory Encoding. J Exp Psychol Learn Mem Cogn 34: 791–808, 2009.

11. Fisk JE, Warr PB. Associative learning and short-term forgetting as a function of age, perceptual speed, and central executive functioning. J Gerontol Psychol Sci 53: 112–121, 1998.

12. Fu Q, Hasan Z, Santello M. Transfer of learned manipulation following changes in degrees of freedom. J Neurosci 31: 13576–84, 2011.

13. Fu Q, Zhang W, Santello M. Anticipatory planning and control of grasp positions and forces for dexterous two-digit manipulation. J Neurosci 30: 9117–26, 2010.

14. Gagnon H, Simmonite M, Cassady K, Chamberlain J, Freiburger E, Lalwani P, Kelley S, Foerster B, Park DC, Petrou M, Seidler RD, Taylor SF, Weissman DH, Polk TA. Michigan Neural Distinctiveness (MiND) study protocol: Investigating the scope, causes, and consequences of age-related neural dedifferentiation. BMC Neurol 19: 1–17, 2019.

15. Gallen CL, Baniqued PL, Chapman SB, Aslan S, Keebler M, Didehbani N, Esposito MD. Modular Brain Network Organization Predicts Response to Cognitive Training in Older Adults. PLoS One 11: 1–17, 2016.

16. Gordon AM, Forssberg H, Johansson RS, Westling G. The integration of haptically acquired size information in the programming of precision grip. Exp brain Res 83: 483–8, 1991.

17. Hess TM. Memory and aging in context. Psychol Bull 131: 383–406, 2005.

18. Heuninckx S, Wenderoth N, Swinnen SP. Systems neuroplasticity in the aging brain: Recruiting additional neural resources for successful motor performance in elderly persons. J Neurosci 28: 91–99, 2008.

19. Hirsiger S, Koppelmans V, Mérillat S, Liem F, Erdeniz B, Seidler RD, Jäncke L. Structural and functional connectivity in healthy aging: Associations for cognition and motor behavior. Hum Brain Mapp 37: 855–867, 2016.

20. Ho J, Tumkaya T, Aryal S, Choi H, Claridge-Chang A. Moving beyond P values: data analysis with estimation graphics. Nat Methods 16: 565–566, 2019.

21. Hutchinson S, Kobayashi M, Horkan CM, Pascual-Leone A, Alexander MP, Schlaug G. Age-related differences in movement representation. Neuroimage 17: 1720–1728, 2002.

22. Jernigan T, Archibald S, Fennema-Notestine, C Gamst A, Stout J, Bonner J, Hesselink J. Effects of age on tissues and regions of the cerebrum and cerebellum. Neurobiol Aging 22: 581–594, 2001.

23. Lukos JR, Ansuini C, Santello M. Choice of contact points during multidigit grasping: effect of predictability of object center of mass location. J Neurosci 27: 3894–3903, 2007.

24. Lukos JR, Ansuini C, Santello M. Anticipatory control of grasping: independence of sensorimotor memories for kinematics and kinetics. J Neurosci 28: 12765–12774, 2008.

25. Mojtahedi K, Fu Q, Santello M. Extraction of Time and Frequency Features from Grip Force Rates during Dexterous Manipulation. IEEE Trans Biomed Eng 62: 1363–75, 2015.

26. Mutter SA, Holder JM, Mashburn CA, Luna CM. Aging and the role of attention in associative learning. Psychol Aging 34: 215–227, 2019.

27. Nowak DA, Berner J, Herrnberger B, Kammer T, Gron G, Schonfeldt-Lecuona C. Continuous theta-burst stimulation over the dorsal premotor cortex interferes with associative learning during object lifting. Cortex 45: 473–482, 2009.

28. Nowak DA, Koupan C, Hermsdörfer J. Formation and decay of sensorimotor and associative memory in object lifting. Eur J Appl Physiol 100: 719–726, 2007.

29. Oliviero A, Profice P, Tonali PA, Pilato F, Saturno E, Dileone M, Ranieri F, Di Lazzaro V. Effects of aging on motor cortex excitability. Neurosci Res 55: 74–77, 2006.

30. Parikh PJ, Cole KJ. Handling objects in old age: Forces and moments acting on the object. J Appl Physiol 112: 1095–1104, 2012.

31. Parikh PJ, Fine JM, Santello M. Dexterous Object Manipulation Requires Context-Dependent Sensorimotor Cortical Interactions in Humans. Cereb Cortex 30: 3087–3101, 2020.

32. Parikh PJ, Santello M. Role of human premotor dorsal region in learning a conditional visuomotor task. J Neurophysiol 117: 445–456, 2017.

33. Pitcher JB, Ogston KM, Miles TS. Age and sex differences in human motor cortex input-output characteristics. J Physiol 546: 605–613, 2002.

34. Rao N, Chen YT, Ramirez R, Tran J, Li S, Parikh PJ. Time-course of pain threshold after continuous theta burst stimulation of primary somatosensory cortex in pain-free subjects. Neurosci Lett 722: 134760, 2020.

35. Rao N, Parikh PJ. Fluctuations in Human Corticospinal Activity Prior to Grasp. Front Syst Neurosci 13: 1–15, 2019.

36. Salimi I, Frazier W, Reilmann R, Gordon AM. Selective use of visual information signaling objects’ center of mass for anticipatory control of manipulative fingertip forces. Exp brain Res 150: 9–18, 2003.

37. Salthouse TA. A theory of cognitive aging. North-Holland, 1985.

38. Seidler RD, Bernard JA, Burutolu TB, Fling BW, Gordon MT, Gwin JT, Kwak Y, Lipps DB. Motor control and Aging: Links to age-related brain structural, functional and biomechanical effects. Neurosci Biobehav Rev 34: 721–733, 2011.

39. Small SA, Stern Y, Tang M, Mayeux R. Selective decline in memory function among healthy elderly. Neurology 52: 1392–1392, 1999.

40. Stevens WD, Hasher L, Chiew KS, Grady CL. A neural mechanism underlying memory failure in older adults. J Neurosci 28: 12820–12824, 2008.

41. Vakil E, Agmon-Ashkenazi D. Baseline performance and learning rate of procedural and declarative memory tasks: younger versus older adults. J Gerontol B Psychol Sci Soc Sci 52: P229–34, 1997.

42. Woodruff-Pak DS, Vogel RW, Ewers M, Coffey J, Boyko OB, Lemieux SK. MRI-Assessed Volume of Cerebellum Correlates with Associative Learning. Neurobiol Learn Mem 76: 342–357, 2001.

43. Zach N, Inbar D, Grinvald Y, Bergman H, Vaadia E. Emergence of novel representations in primary motor cortex and premotor neurons during associative learning. J Neurosci 28: 9545–9556, 2008.

44. Zhang W, Gordon AM, Fu Q, Santello M. Manipulation after object rotation reveals independent sensorimotor memory representations of digit positions and forces. J. Neurophysiol. (2010). doi: 10.1152/jn.00140.2010.

45. Zhong JY, Moffat SD. Age-related differences in associative learning of landmarks and heading directions in a virtual navigation task. Front Aging Neurosci 8: 1–11, 2016.

